# DYRK1A role in microtubule-based axonal transport regulates the retrograde dynamics of APP vesicles in human neurons

**DOI:** 10.1101/2021.02.25.432601

**Authors:** Iván Fernandez Bessone, Karina Karmirian, Livia Goto-Silva, Mariana Holubiec, Jordi L. Navarro, Emanuel Martínez, Trinidad M.M. Saez, Stevens Rehen, Tomás L. Falzone

**Author notes:** Correspondence addressed to T.L.F.

## Abstract

In Alzheimer’s Disease (AD) the abnormal intracellular distribution of the amyloid precursor protein (APP) affects its processing and, consequently, the generation of Aβ. Axonal transport plays key roles in the neuronal distribution of APP. The dual-specificity-tyrosine phosphorylation-regulated-kinase-1A (DYRK1A) has been associated with AD onset since its overexpression was found in Down syndrome and sporadic AD patients. Experimental evidence confirmed that APP and tau phosphorylations are mediated by DYRK1A. Moreover, DYRK1A can regulate the cytoskeletal architecture by phosphorylation of both tubulin subunits and microtubule-associated proteins. Therefore, we tested whether DYRK1A has a role in APP axonal transport regulation.

We developed highly-polarized human-derived neurons in 2D cultures. At day 14 after terminal plating we inhibited DYRK1A for 48hs with harmine (7.5 μM). DYRK1A overexpression was induced to perform live-cell imaging of APP-loaded vesicles in axons and analyzed transport dynamics. A custom-made MATLAB routine was developed to track and analyze single particle dynamics.

Short-term harmine treatment reduced axonal APP vesicles density, due to a reduction in retrograde particles. Contrarily, DYRK1A overexpression enhanced axonal APP density, due to an increase in the retrograde and stationary component. Moreover, both harmine-mediated DYRK1A inhibition and DYRK1A overexpression revealed opposite phenotypes on single particle dynamics, affecting primarily dynein processivity. These results revealed an increased retrieval of distal APP vesicles in axons when DYRK1A is overexpressed and reinforce the suggestion that DYRK1A enhance APP endocytosis‥

Taken together our results suggest that DYRK1A has a relevant role in the regulation of axonal transport and sub-cellular positioning of APP vesicles. Therefore, our work shed light on the role of DYRK1A in axonal transport regulation, and the putative use of harmine to restore axonal transport impairments.

## Introduction

In Alzheimer Disease (AD) the abnormal metabolism of the amyloid precursor protein (APP) is linked to impairments in axonal transport primarily affecting the intracellular distribution of APP and the extent to which is exposed to secretases that generate amyloid beta peptides (Aβ)^1 2 3^. Distinct kinases regulate the distribution of APP within neurons by the specific phosphorylation of its intracellular domain or by modifying proteins that regulate its transport^4^. Recently, a reduction of phosphorylated APP was described after treating neurons with harmine, a β-carboline alkaloid with high potency to inhibit DYRK1A ^5 6 7^. Moreover, harmine-mediated DYRK1A inhibition improved spatial learning and memory in AD mouse models, raising it as a putative therapeutical strategy. *DYRK1A* gene is located within the critical region on chromosome 21 that is triplicated in Down syndrome (DS) implying a synergistic effect between APP and DYRK1A in the early progression of disease and the manifestation of AD pathology^8 9 10 11^. In addition, increased DYRK1A levels were found in postmortem analysis from sporadic AD brains^12^. However, the role of DYRK1A in APP neuronal distribution and/or in the regulation of axonal transport mechanism in human neurons is not known.

The early temporal and spatial expression of *DYRK1A* in neurons indicate a role in the central nervous system development ^13 14 15^. Relevant roles are also played by DYRK1A in the maintenance of adult brain functions^16 17^. *DYRK1A* genetic deletion in fly and mouse models or the loss of function mutations in humans lead to severe neuronal developmental disorders including microcephaly^18 19 20 21 22^. On the other hand, *DYRK1A* triplication in DS together with the *APP* gene, also located in the critical region of chromosome 21, support a genetic interaction that enhance the neurodegenerative phenotype. Even microduplication of chromosome 21 including *DYRK1A* and not *APP* have been implicated to impair neuronal function and lead to DS ^23^. Several fold increases in the number of DYRK1A-positive neurofibrillary tangles (NFTs) were observed in DS and AD brains suggesting a direct contribution of DYRK1A to the acceleration and enhancement of the neuronal loss observed in disease ^24^.Due to its nuclear import signal DYRK1A has been described as a key regulator of tau gene (*MAPT)* expression by the specific phosphorylation of the alternative splicing factor (ASF) that reduces its ability to promote tau exon 10 inclusion in DS ^25^. However, subcellular fractionation-based and confocal microscopy studies revealed the majority (almost 80%) of DYRK1A in the cytoplasm of neurons, supporting a role in the direct regulation of cytosolic proteins ^14 16 24^. Cytoplasmic DYRK1A mediates the phosphorylation of AD related proteins such as APP and tau (cita). APP phosphorylation at Thr668 is associated with increased Aβ generation in the brains of transgenic mice that overexpress the human DYRK1A protein^26^. DYRK1A also phosphorylates tau at multiple sites in several cellular models and act as a priming kinase for further tau phosphorylation by glycogen synthase kinase 3 (GSK3) ^27 28^. Many processes regulated by DYRK1A contributes to the intracellular accumulation of abnormal cytoskeleton proteins, including the formation of NFTs^17 24 25^ and Lewy bodies^12 26^. Interestingly, the reduction of DYRK1A activity has emerged as a relevant target for the treatment of neurodegenerative diseases due to its functional activity on multiple pathways implicated in neuronal function and AD^12^. Harmine, selectively inhibit DYRK1A activity and reduce tau phosphorylation at multiple disease-related sites ^8^. This specificity is confirmed since other inhibitors of DYRK1A also reduced tau pathology^29^. Since the chronic inhibition of DYRK1A reduces insoluble forms of Aβ and hyperphosphorylated tau, its long-term use is associated with delay in the onset of both amyloid plaques and NFTs in mouse models ^30^. The diversity of DYRK1A substrates and their localization in different cell compartments have supported new functions for this kinase in regulating intracellular trafficking mechanisms ^31^. Consistent with the detection of DYRK1A in synaptosomes, it was proposed a novel role in neuronal endocytosis since it can phosphorylate dynamin 1, amphiphysin 1, and synaptojanin 1; a group of proteins involved in synaptic vesicle recycling ^32 33 34 35 36^. These findings suggest that DYRK1A regulate multiple steps in the formation and uncoating of clathrin-coated vesicles^36^. Moreover, other functions of DYRK1A were posed over neuronal polarity regulation and maintenance since through phosphorylation can control the binding of proteins to microtubules. The growing list of DYRK1A cytoskeletal targets including tau, α-synuclein, SEPT4, MAP1B and N-WASP ^28 37 38 39 40^, support a significant role in the regulation of cytoskeleton structure associated with intracellular dynamics that has not been studied in detail.

Axonal transport is a complex multistep process highly regulated by the action of different kinases that ensure the stabilization and maintenance of neuronal polarization and function. APP, as a type I transmembrane protein, is packed in Golgi-derived vesicles that are massively distributed to different locations in neurons^41 42 43^. The axonal transport of APP is necessary to maintain synaptic boutons, promote axonal growth as a guidance molecule, and act as a rapid axonal injury signal^44 45 46 47^. Changes in APP transport are linked to its processing and vice versa since increased β-secretase cleavage reduces the anterograde delivery of APP to synapses, while reduced APP axonal transport and vesicle density are induced when APP is re-routed to the endocytic pathway for processing after proteasome inhibition or selective UV irradiation^48 49 50 51^. It has been proposed that phosphorylation of APP controls its neurite localization due to a transport dependent mechanism ^52 53^, and this process either by phosphorylation or redistribution leads to enhanced APP processing ^4^. A wide variety of kinases including GSK3, MAP kinases, cyclin-dependent protein kinase 5 (cdk5), casein kinase 1 and 2 (CK1, 2), MAP/microtubule affinity-regulating kinases (MARKs), have been proven to regulate axonal transport since can phosphorylate one or more regulatory proteins such as microtubules, microtubules associated proteins, motors and/ or cargos ^54^. DYRK1A has shown to act on tau, APP, microtubules associated proteins and proteins involved in membrane recycling, however, still remain unknown whether DYRK1A can act as a direct regulator of APP axonal transport.

Here we test the hypothesis that DYRK1A activity impact in the process of intracellular dynamics that are key for polarized neuronal function and specifically in the regulation of the axonal transport of the APP vesicle. Our results highlight DYRK1A as a new player in the complex microtubule-based transport pathway and bring light into the metabolism of APP through the synergistic interaction proposed between APP and DYRK1A, which has relevance for the progression of neurodegenerative diseases such as AD.

## Results

### Harmine regulates the axonal transport of APP by reducing its retrograde component

We proposed to study the impact of harmine on the axonal transport of the APP vesicle in highly polarized human-derived neurons obtained following a protocol described previously^55^ (Materials and Methods, Supplementary 1). Fourteen days-old human enriched neuronal cultures express neuronal markers (βIII-tubulin, green) with few cells expressing neural precursor marker nestin (red)(Fig 1A). Moreover, phosphorylated tau and ankyrin-G staining was observed in neuronal projections and at the axon initial segment (AIS), respectively, revealing the presence of mature axonal structures associated with the generation of action potentials (Fig 1B). These cultures were used to test the effect of harmine on microtubule-based transport by performing high resolution/speed live-cell imaging to record the axonal transport of APP vesicles. Human neurons were transfected with the pcDNA-APP-YFP vector driving the expression of the yellow fluorescent protein-tagged APP at the C-terminal region (APP-YFP) at day 14 after terminal plating (D14) (Fig 1C). Transfected neurons showed a vesicular distribution of APP-YFP along the axon (Fig. 1B). Human neurons treated either with vehicle (DMSO) or with harmine at 7.5 μM to inhibit DYRK1A for 48 hrs showed no morphological or structural changes and maintain unchanged the AIS cytoskeletal structure (Fig 1E). APP-YFP axonal transport was registered in axons from DMSO and harmine treated cultures and 30 seconds movies were transformed to kymographs from which trajectories were tracked semi-automatically using custom-made MATLAB routines (Figure 1F). MATLAB algorithms were used to extract axonal transport properties revealing that harmine induced a significant decrease in the density of axonal APP vesicles (Fig 1G). Interestingly, these changes were observed due to a selective reduction in the density of APP retrograde moving vesicles without changing the anterograde or stationary vesicle density in neurons treated with harmine compared with control (Fig. 1H). Since the load of retrograde vesicles into the axon was changed by harmine we ask whether DYRK1A inhibition can also regulate the processive movement of APP in axons. Therefore, we analyzed single particle trajectories of moving APP-YFP vesicles using a modification of previously validated MATLAB routines ^56 57^. In-depth analysis of transport dynamics revealed that harmine induce a significant increase in the number of anterograde and retrograde runs only in the net retrograde moving vesicles without impairing the anterograde ones (Fig 1I, J). Contrarily, the distance and duration of independent runs was not affected after harmine treatment (Fig 1K, L). Together, these results suggest that harmine exert changes in APP axonal transport that are selectively associated with reductions of retrograde vesicle density load into axons and impairments in the retrograde number of runs associated with dynein processivity and retrograde motor activity.

**Figure 1.**
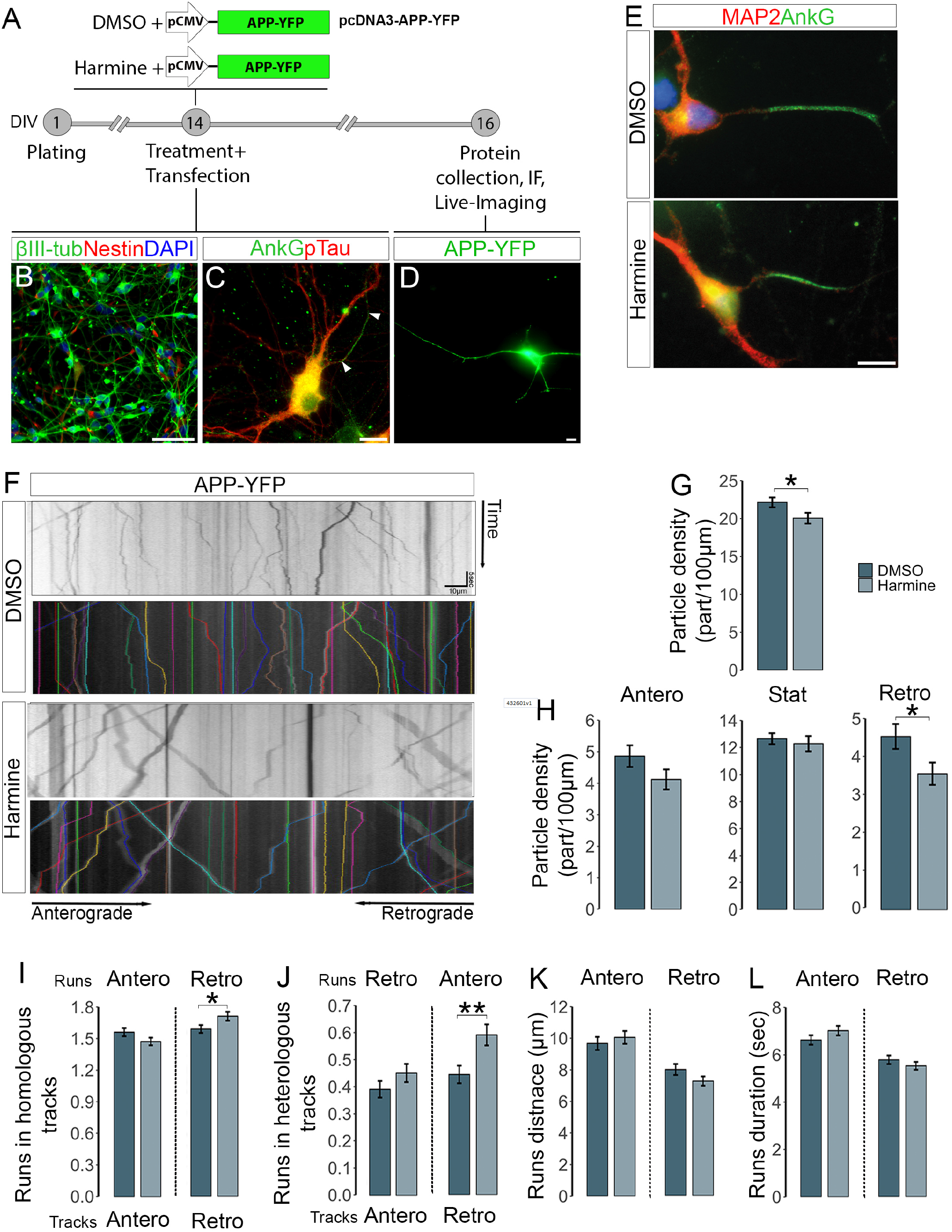
Harmine regulates the axonal transport of APP by reducing its retrograde component‥. (A) Schematic representation of experimental design used to analyze APP dynamics in control (DMSO) or harmine (7,5 μM) in human polarized neurons in 2D‥ (B-D) Confocal images of 2D culturesenriched in highly polarized human neurons. (B) Immunofluorescent staining against the neuronal marker βIII tubulin (red) and the neural precursor marker Nestin (green) in 14 days-old neurons after terminal plating (D14). (C) Immunofluorescent staining against the neuronal AIS marker AnkG (green) and phosphorylated tau (red). (D) 16D human neuron transfected with pcDNA-CMV-APP-YFP (APP-YFP) and immunostained against the green fluorescent protein (GFP). (E) Immunofluorescent staining against AnkG after 48 hrs of DMSO or harmine (7,5 μM) treatment in D16 human neurons. DAPI was used for nuclear staining (blue). Scale bar 50 μm (B), 10 μm (C, D, E). (F) Kymographs of time versus distance obtained from a 30s movie (8 frames/seconds) recorded in axons from neurons transfected with APP-YFP and treated with DMSO or harmine (7.5 μM) for 48hs. Colored lines represent trajectories recovered from a semi-automatic tracking tool box system. (G-H) Average APP vesicle density in axons from control (DMSO) and harmine treated neurons. Total (G), anterograde, stationary, and retrograde (H) APP vesicle density in 100 μm axon length. Kymographs n = 98 DMSO and 116 harmine from 3 independent experiments. Each kymograph represents an individual neuron. (J, I) Average number of homologous (I) and heterologous (J) runs in anterograde and retrograde moving APP vesicles. Anterograde vesicles n=570 DMSO, n=651 harmine; retrograde vesicles n=548 DMSO, n=553harmine.Average run lengths distance (K) and duration (L) from APP vesicles travelling in a net anterograde or retrograde direction. Anterograde runs n=1287 DMSO, n=1475 harmine; retrograde runs n=1247 DMSO, 1431 harmine.Data presented as mean ± s.e.m. Student’s t-test *P<0.05, ***P<0.01, ****P<0.001.

### DYRK1A overexpression enhance the retrograde component of APP axonal transport

Genomic *DYRK1A* triplication in DS and its overexpression in sporadic AD suggest that increased DYRK1A activity induce a detrimental effect in neuronal function that leads to neurodegeneration^58^. To unravel the effect of excess DYRK1A in APP axonal transport without disturbing its early role in neural differentiation and polarization ^15^ we induced DYRK1A overexpression for 48 hs in already polarized human neuronal cultures at D14 (Fig 2A). A vector driving DYRK1A expression fused to the mCherry fluorescent marker (DYRK1A-mCherry) was subcloned under the CMV promoter and its expression was confirmed in western blots from homogenates of lipofectamine transfected N2A neuroblastoma cell lines (Fig 2B). At D16, human neurons co-transfected with APP-YFP (green) and DYRK1A-mCherry (red) showed a significant (100%) increase in DYRK1A fluorescent intensity when compared with non-transfected neurons (Fig 2C). Using this setting we then analyzed the axonal transport of APP in control (APP-YFP+mCherry) and DYRK1A overexpression (APP-YFP+DYRK1A-mCherry). Thirty seconds movies registering the transport of APP vesicles within axons were transformed to kymographs from which trajectories were semi-automatically tracked using custom-made MATLAB routines (Fig 2D). Interestingly, DYRK1A overexpression induce a significant increase (25%) in total axonal APP vesicle density, contrarily to harmine effect (Fig 2E). Moreover, selective changes were observed by an increase in stationary and retrograde densities of APP vesicles again without anterograde impairments (Fig 2F). In-depth analysis of transport dynamics revealed no changes in the number of homologous or heterologous runs both in anterograde and retrograde net moving vesicles (Fig 2H). However, higher retrograde processivity was observed after DYRK1A overexpression by an increase in the distance travelled for retrograde runs (Fig 2I) without changes in runs duration (Fig 2J). These results showed an increased retrieval of distal APP vesicles in axons when DYRK1A is overexpressed and reinforce the suggestion that DYRK1A enhance APP endocytosis and selectively modulate the retrograde component of APP by increasing the processivity of dynein motors.

**Figure 2.**
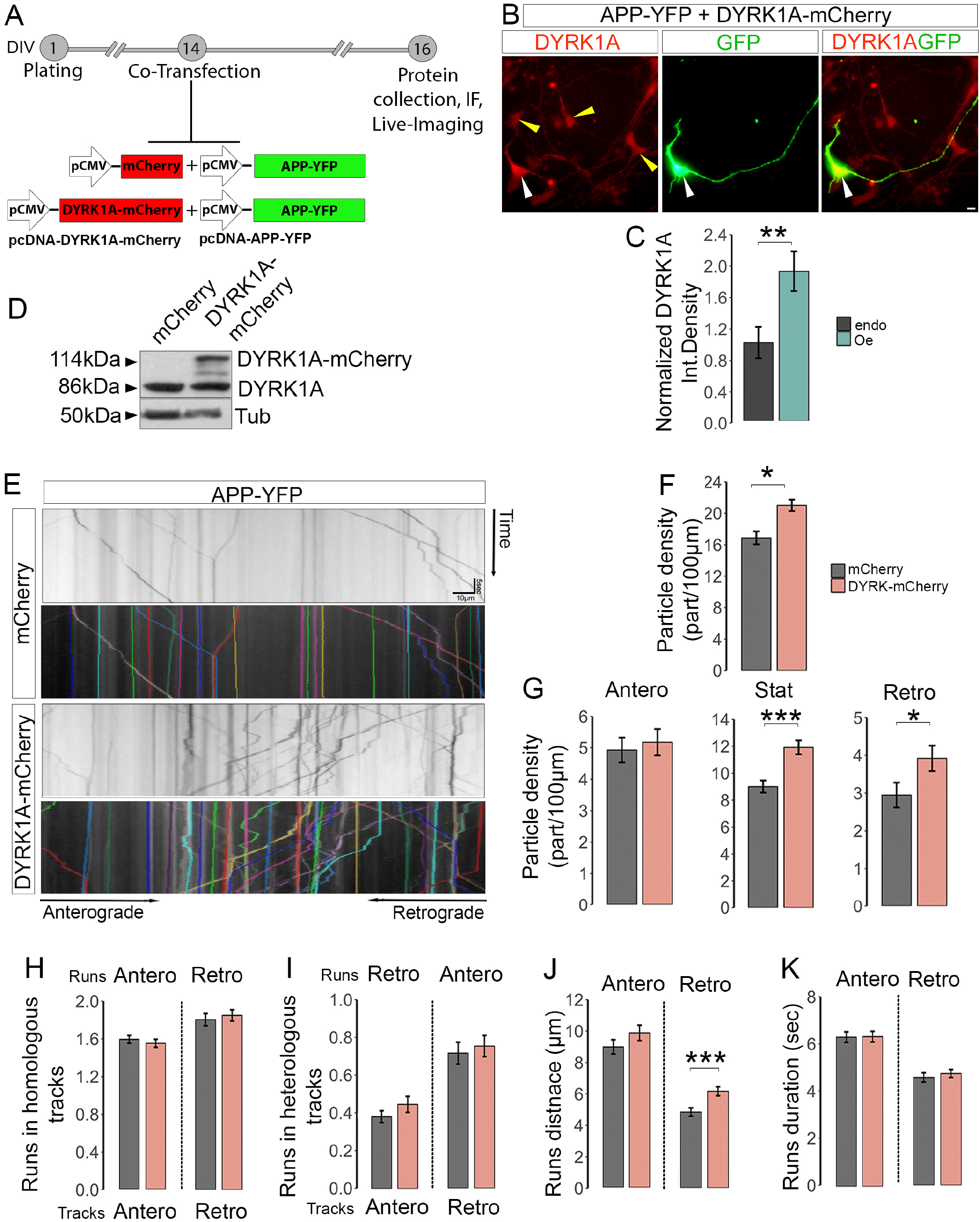
DYRK1A overexpression enhance the retrograde axonal transport of APP‥. (A) Schematic representation of experimental design used to analyze the effect of DYRK1A over-expression on APP-YFP axonal transport dynamics in D16 human neurons. (B) Human neuronal culture at D16 co-transfected with pcDNA-CMV-DYRK1A-mChrerry (DYRK1A-mChrerry, red) and APP-YFP (green) for 48 hours and immunostained with DYRK1A and GFP antibodies (white arrow). Scale bar 10 μm. (C) Quantification of fluorescent integrated density of DYRK1A in co-transfected (white arrow) neurons normalized to endogenous levels in non-transfected (yellow arrows) neurons. (D) Western blots showing endogenous DYRK1A (86 kDa) and transfected DYRK1A-mCherry (116 kDa) protein levels from control (mCherry) or DYRK1A-mChrerry transfected Neuro2A homogenates. (E) Kymographs of time versus distance obtained from a 30 s movie (8 frames/seconds) recorded in axons from neurons co-transfected with APP-YFP+mCherry (top) or APP-YFP+DYRK1A-mCherry (bottom). Colored lines represent trajectories recovered from a semi-automatic tracking tool box system. (F, G) Average APP vesicle density in axons from control (APP-YFP+mCherry) and DYRK1A over-expressing (APP-YFP+DYRK-mCherry) neurons. Total (F), anterograde, stationary, and retrograde (G) APP vesicle density in 100 μm axon length. Kymographs n = 62 APP-YFP+mCherry, n=54 APP-YFP+DYRK-mCherry from 3 independent experiments. Each kymograph represents an individual neuron. (H, I) Average number of homologous (H) and heterologous (I) runs in anterograde and retrograde moving APP vesicles. Anterograde vesicles n=570 APP-YFP+mCherry, n=651APP-YFP+DYRK-mCherry; retrograde vesicles n=548 APP-YFP+mCherry, n=553APP-YFP+DYRK-mCherry. Average run lengths distance (J) and duration (K) from APP vesicles travelling in a net anterograde or retrograde direction. Anterograde runs n=1287 APP-YFP+mCherry, n=1475 APP-YFP+DYRK-mCherry; retrograde runs n=1247 APP-YFP+mCherry, n=1431APP-YFP+DYRK-mCherry. Data presented as mean ± s.e.m. Student’s t-test *P<0.05, ***P<0.01, ****P<0.001‥

### Harmine and DYRK1A overexpression revealed a selective modulation in retrograde molecular motor mechanisms of APP axonal transport

The *in vivo* transport of APP is proposed to occur due to a concerted action of several motors in a cooperative multi-motor arrangement. The probability of transitions between velocities in a multi-motor system can increase or decrease depending on liability exerted over motor number exchange. To study whether DYRK1A inhibition or overexpression modulate molecular motors configuration within APP loaded vesicles,we ploted heat-map graphs from the probability of a given velocity to change to a slower or higher next velocity in the anterograde or retrograde condition (Figure 3 A-F). This allowed the identification of higher probability of transitions between anterograde to anterograde runs (Q1) and retrograde to retrograde runs (Q3) than those observed for reversions (Q2, Q4) (Figure 3C). To compare both inhibition and overexpression conditions, the probability difference between harmine and DMSO or between DYRK1A and mcherry were computed using the sum of absolute values analyses to find almost no changes in velocity transition in both experimental conditions. These results that either increased or decreased DYRK1A activity does not modify the probability of exchanging molecular motor arrangements in the anterograde and retrograde direction. Since kinesin and dynein forces interact in a tag of war/coordination model to exert directional transport in axonal polarized microtubules^59^, we ask whether harmine or DYRK1A overexpression modifies the segmental velocities of APP cargo that are directly correlated with the number of active motors in a given condition^60^. Therefore, segmental velocities were extracted from tracked APP vesicles registered in kymographs obtained from previous experimental settings. A multimodal distribution of segmental velocities was observed, as previously described ^57^, so we propose a Gaussian mixture model with three components to describe each dataset (Materials and Methods). Segmental velocity distribution of moving APP vesicles obtained under harmine treatment showed similar anterograde velocity modes (A, B, C) when compared with DMSO treated neurons (Fig. 3G, Supplementary 2). However, mild shift towards faster retrograde velocities were observed in modes B and C when plotting retrograde distributions (Fig 3H, Supplementary 2). Interestingly, after DYRK1A overexpression velocity mode distributions for the anterograde APP moving vesicles were again similar to control (Fig 3I, Supplementary 2). Nevertheless, significant increases were observed towards faster retrograde velocity mode distribution in modes A, B and C after DYRK1A overexpression compared with control (mCherry) (Fig 3J, Supplementary 2). Together, these results strengthen the suggestion that DYRK1A activity induce a selective modulation of the retrograde component in APP vesicle transport by changing the number of active dynein motors associated with the distribution of retrograde velocities.

**Figure 3.**
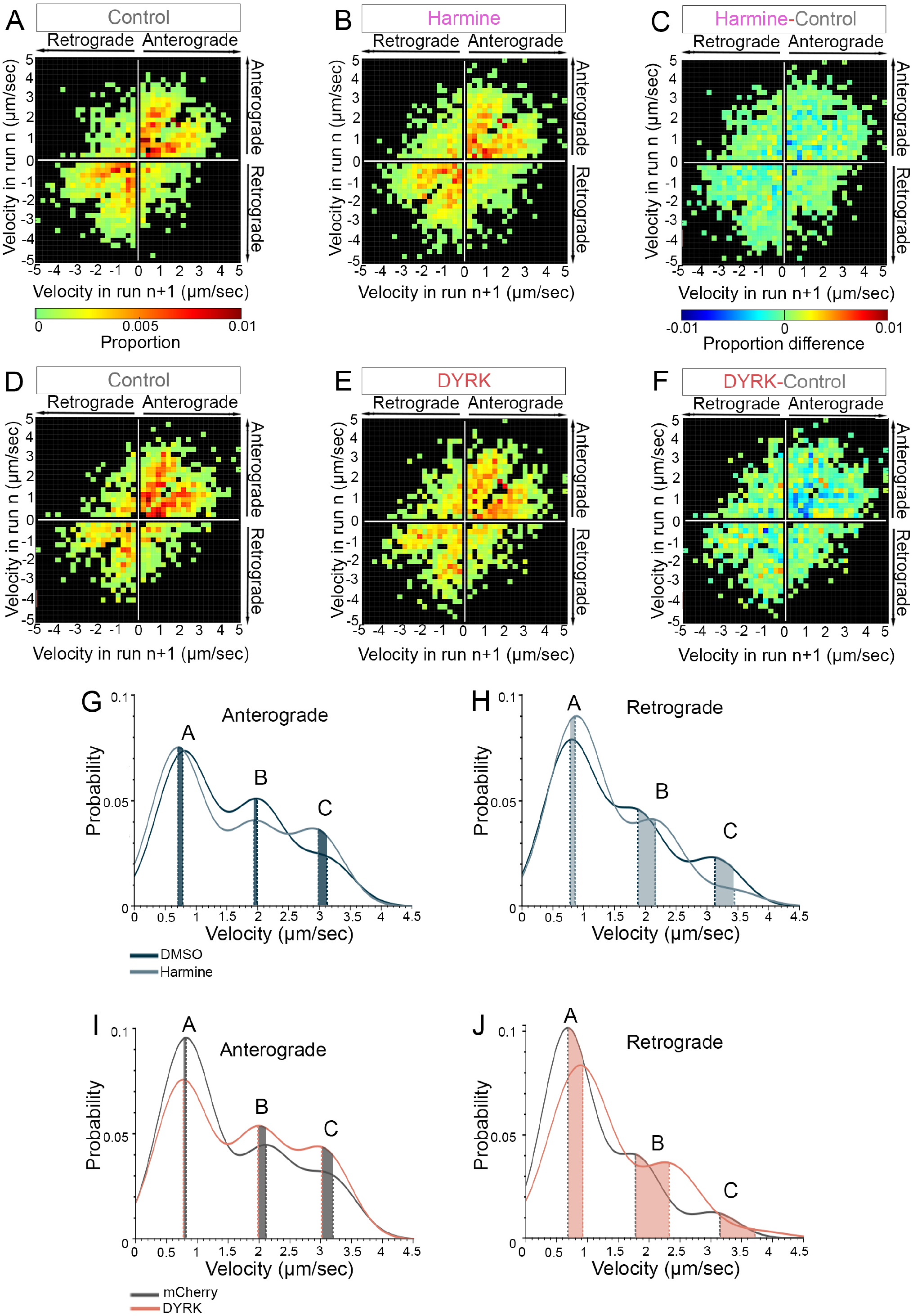
Harmine and DYRK1A over-expression revealed a selective modulation in retrograde molecular motor mechanisms of APP axonal transport. (A-C) Heat maps showing the probability of velocity transitions from velocity n to velocity n+1 in control (A, DMSO) and after harmine treatment (B). (C) Heat map showing the positive (red) or negative (blue) frequency from the difference between experimental (harmine) minus control (DMSO). (D-F) Heat maps showing the probability of velocity transitions from velocity n to velocity n+1 in control (D, mCHerry) and after DYRK1A overexpression (E, DYRK1A-mCherry). (F) Heat map showing the positive (red) or negative (blue) frequency from the difference between experimental (DYRK1A-mCherry) minus control (mCherry). Non significant differences obtained from comparison of sum of absoluts between controls and simulations, and control and experimental conditions per quadrant. The sum of absoluts between simulations and control and control and experimental are displayed in Table 1(G-J) Distribution of segmental velocities of moving APP–YFP vesicles represented as a Gaussian mixture model with three modes (A–C) associated with number of active motors. Anterograde (G) and retrograde (H) segmental velocities distribution of APP vesicle transport in DMSO (dark blue) and harmine-treated (light blue) human neurons. Anterograde (I) and retrograde (J) segmental velocities distribution of APP vesicle transport in mCherry (dark red) and DYRK-mCherry (light red) human neurons. Mode center represented by dotted line. The center and fraction of the different modes are displayed in Table 1. Segmental velocities n= 7694 anterograde and 6412 retrograde DMSO; 9225 anterograde and 7177 retrograde harmine; 5954 anterograde and 3177 retrograde mCherry; 5788 anterograde and 3762 retrograde DYRK-mCherry. Significant differences (**) obtained from comparison of non-overlapping confidence intervals with each control group.

## Discussion

Here, using human functional human neurons derived from induced pluripotent stem cells (iPSC), we searched for neuronal processes modulated by harmine-mediated DYRK1A inhibition. We performed high-definition/high-speed live imaging of APP vesicles in 2D human-derived neurons after modulating DYRK1A activity to find specific changes in the regulation of retrograde APP axonal vesicle loading and in dynein-dependent motor protein transport. Our experiments identified new roles for DYRK1A in the control of intracellular dynamics and specific changes in the retrieval of APP towards the cell body. Together, these findings highlight DYRK1A inhibition as a putative therapeutic target to restore APP metabolism and axonal transport defects in neurodegenerative diseases such as AD.

DYRK1A inhibition has been proposed as a putative therapeutic strategy in AD, since it reduces APP phosphorylation and has a positive impact on restoring abnormal phenotypes in AD mouse models^7 30 61 62 63^. To study the interaction of DYRK1A and APP we cultured human differentiated neurons to analyze the axonal transport properties of fluorescently labeled APP vesicles after DYRK1A modulation. Interestingly, harmine significantly reduced APP axonal density by a selective decrease in the retrograde component, suggesting that APP vesicle loading at the axonal tip was reduced by DYRK1A inhibition. Noteworthy, DYRK1A regulates the assembly of the endocytic apparatus, by phosphorylating multiple endocytic proteins required for clathrin-coated vesicle (CCV) formation^35 36^. APP is found within newly formed endocytic CCV, suggesting a putative regulation of APP internalization by DYRK1A ^64 65^.These results were reinforced since DYRK1A overexpression increased APP axonal density by the selective enhancement of retrograde and stationary vesicles. Interestingly, phosphorylation of dynamin by DYRK1A appears to have a dual role in regulating the interaction of dynamin with amphiphysin and endophilin, a key step in synaptic vesicle internalization ^66^. Experimental evidence suggests that DYRK1A overexpression delay the initial process of internalization by reducing CCV assembly^67 68^. However, this dual role in mediating protein binding for CCV formation plus the observation that DYRK1A activity is necessary for the uncoating of endocytosed CCV may result in complex mechanisms necessary for the axonal loading and release of APP into transport^69^. Moreover, the observed DYRK1A mediated phosphotylation of APP at Thr668 can be accounted as another regulatory step for APP internalization and colocalization with early endosomes ^26 70^. APP cleavage into the pathological amyloidogenic form occurs mainly in intracellular endolysosomal compartments^3 50^. Therefore, enhanced retrograde density due to DYRK1A overexpression may provide a critical pool of APP to the endo-lysosomal system for β-secretase processing^71 72^. On the other hand, harmine reduced the density of retrograde APP, suggesting that DYRK1A inhibition may delay it amyloidogenic processing, and support the therapeutic approach directed to delay the onset of amyloid - like pathology in AD ^30^.

From many targets, DYRK1A also phosphorylate different tubulin subunits and microtubule associated proteins affecting microtubule architecture ^39 73^. Axonal transport depends on microtubule stability that is highly regulated by protein post-translational modifications ^74^. However, the effect of DYRK1A on microtubule-mediated transport has not been directly studied. We therefore analyzed DYRK1A-mediated single particle dynamics of axonal APP vesicles to unravel it direct control on microtubule-motor dependent transport. Harmine-treatment reduced the processivity of retrograde APP by inducing stochastic movements that increase the number of runs in retrograde trajectories. Contrarily, DYRK1A overexpression facilitated the retrograde processivity by increasing the distance travelled of retrograde runs. Numerous evidence suggests that microtubule associated protein composition is essential for recruiting or repelling molecular motors to the axonal microtubule lattice^75 57^, while other microtubule binding proteins act as molecular brakes for retrograde transport in dendrites ^76^. Our results suggest that DYRK1A-mediated modifications either modify the MAP code and/or change the molecular motor composition within APP vesicles leading to specific changes in retrograde APP transport ^57^. Contrary to the possibility of continuous exchanges in motor configuration, we observed no significant variations in the probability of velocity transitions after DYRK1A modulation, suggesting that DYRK1A is not affecting the on/off state of molecular motors. Noteworthy, DYRK1A overexpression revealed a significant increase in the retrograde segmental velocity modes that is associated with an increased number of vesicles with loaded active motors. Interestingly, upon an obstacle encounter, cargoes transported by dynein-dynactins often remain attached to the microtubule and undergo microtubule sliding^77^. Moreover, differential effects are exerted on kinesin and dynein motors by microtubule protein decoration^57 78 79^. DYRK1A appears to selectively modify the retrograde component due to changes in microtubule decoration and/or the regulation of active dynein motors in distal axonal loaded vesicles. Future effort should be directed to dissect the effect of DYRK1A on individual proteins that either regulate vesicle loading, microtubule protein decoration, and dynein motor configurations in APP transport.

Our work highlights DYRK1A as a modulator of microtubule associated proteins, motors and adaptors that in combination may play a synergistic role in the regulation of retrograde axonal transport of APP. Moreover, we validated harmine effect in human neural tissue as a modulator of relevant neuronal processes that regulate intracellular dynamics. These results stress the relevance of DYRK1A in the regulation of APP metabolism in neurodegenerative diseases and shed light on a novel molecular pathway that can be targeted for therapeutic interventions in Alzheimer Disease.

## Materials and methods

### hiPSCs differentiation to human neurons

Neurons were derived from CV-hiPSCs line (MTA UCSD, San Diego, USA). Differentiation to functional human neurons was performed as previously described (Pozo Devoto et al, 2017). Briefly, optimal size colonies were enzymatically detached and grown in suspension for 48 hours to allow embryoid body (EB) formation. Then, EBs were neuronally induced in suspension for 3 days (NIM: DMEM/F12 (Gibco, 12634-010), 1% N2 supplement (Gibco, 17502-048), 1% MEM-NEAA (Gibco, 11140-050), 280 UI/ml heparin (Sigma, H3393), and 1% Pen-Strep (Gibco, 15140-122). Embryoid bodies were attached to laminin-coated 6-well plates and grown for 7–14 d to allow the formation and maturation of neural tube-like rosettes enriched in neural progenitors. Neural precursors were manually picked and transferred to 25 cm2 flasks in NIM medium supplemented with B27 and ascorbic acid (NIM + 2% B27 supplement (Gibco, 17504-044), 0.1% ascorbic acid (Sigma Aldrich, A1300000)) for up to 1 month, changing media every 2 days. Neural rosettes were dissociated in 1% trypsin (Gibco, 15090-046) and accutase (1:1) (Gibco, A11105) for 8 min at 37°C and later blocked using 0.5 mg/ml trypsin inhibitor (Gibco, R007100) and the suspension centrifuged for 5 min at 1000 rpm. Pellet was washed with DMEM/F12, disggregated to single-cell and resuspended in neural differentiation media (NDM: Neurobasal medium supplemented with 1% N2 supplement (Gibco, 17502-048), 2% B27 supplement (Gibco, 17504-044)). Cells were plated over poly-ornithine (Sigma-Aldrich, P4957) and laminin-coated (Gibco, 23017-015) coverslips (0.1 mg/ml and 20 μg/ml, respectively) into 24-well plates and maintained in 500 μl/well complete NDM medium (NDM + 1 μg/ml laminin (Gibco, 23017-015), 1μM cAMP (Sigma-Aldrich, A6885), 200 μg/mL ascorbic acid (Sigma Aldrich, A1300000), 10 ng/ml BDNF (Gibco,10908-010), 10 ng/ml GDNF (Gibco, PHC7041)). Half of the media was replaced every 3 days.

### Western blotting

Total protein from neurons or cell lineswas collectedin 100 μl of lysis buffer [50 mM Tri-HCl (pH 7.5), 150 mM NaCl, 1% Igepal (Sigma-Aldrich, I8896) and 1x protease inhibitor cocktail (Sigma-Aldrich, P2714)] and centrifuged for 10 min at 10,000 rpm (8000 g) at 4°C. Protein concentration of the supernatant was measured using BCA Protein Assay Kit (Pierce, 23225). Equal amounts of protein (30 μg) in 10 μl of 4x Laemmli sample buffer (250 mM Tri-HCl (pH 6.8), 0.04% bromophenol blue, 40% glycerol, 8% SDS and 2% β-mercaptoethanol) were loaded onto 12% SDS-polyacrylamide gel and See-Plus 2 (Invitrogen, LC5925) was used as a molecular-weight marker. Proteins were transferred onto nitrocellulose membranes using a wet system in 25 mM Tris base, 190 mM glycine and 20% methanol. Membranes were blocked in 5% BSA in 0.1% Tween-20 in TBS (TBS-T) for 1 h and incubated in 1% BSA in TBS-T with primary antibody overnight at 4°C. After washing, membranes were incubated with HRP-conjugated secondary antibody for 4 h at room temperature. Westerns were developed using an ECL kit (Pierce, 32106). Scanned images were analyzed using ImageJ software.

### Immunofluorescence, image collection and axon initial segment morphology

PBS washed cells were fixed (4% paraformaldehyde, 4% sucrose in PBS) for 30 min at 37°C and permeabilized with 0.1% Triton X-100 for 10 min at room temperature (RT). Cells were blocked for 1 h at RT (3% BSA, 0.1% Triton X-100, and 10% goat serum in PBS) and incubated with primary antibodies in blocking solution overnight at 4°C. Secondary antibodies were incubated for 2 h at RT and DAPI (concentracioon) was included in last washed for 30 min.Cells were mounted on slides with DPX (Sigma Aldrich, 06522). AIS morphology was analyzed using ImageJ software.

### Live-cell-imaging

Imaging of live cells and kymograph generation was performed as described previously (Falzone and Stokin, 2012). Briefly, 30 s movies of APP–YFP moving particles in neurons were recorded using an inverted epifluorescence microscope (Olympus IX81) connected to a CCD camera (Olympus DP71/12.5 megapixels). Cultures were observed under a 100Xlens (numerical aperture, NA: 1.25) and maintained at 37°C, 5% CO2, and 10% humidity using a CO2 humid chamber and heated stage (Olympus). Directionality was determined by tracking fluorescent axons. To avoid introducing biases, imaging was performed in axons at their middle part separated by at least two fields of view distance (~200 μm) from cell bodies and from axonal tips. Kymographs were generated from the recordings with ImageJ using the multiple kymograph plug-in (Otero et al., 2014).

### Axonal transport dynamics analysis

Segmental velocities, run lengths, pauses, and reversions were computed from trajectories using custom-made MATLAB routines. For the calculation of segmental velocities, processive trajectories were divided into 20 frames of duration, producing a linear approximation with the least-squares method, filtering trajectories with outliers and significant differences between the dataset and the fitted curve. Subsequently, the slope of the regression was considered as the velocity of the segment. The point in which the movement is equal to 0, filtering the points produced by the noise of the trajectory was used for the calculation of run lengths, pauses, and reversions. The trajectories between those points are considered continuous segments and classified in anterograde or retrograde movement if they show positive or negative velocity respectively (velocities are the slope of a linear approximation of the trajectory). Pauses are considered when segments follow a stationary criteria for >5 frames (625 ms) moving <0.05 pixel per frame (<0.16 μm/s). The script merges continuous segments of the same type and segments that are too small. The difference in micrometers between the final point and the initial point of a determined continuous segment is considered as the run length. The points separating anterograde and retrograde segments are considered reversions. The parameters of the Gaussian mixture model were determined by an expectation maximization algorithm using a function (gmdistribution.fit) in MATLAB. A bootstrapping with resampling procedure with N = 1000 was implemented to compute the intervals of confidence of the Gaussian mixture parameter estimates.

### Statistical analysis

Data processing and statistical analyses were performed using R software.All statistical details of performed experiments, such as performed statistical tests, and valuesof n, are described in the figure legends.t-test was used for comparison of two sample with Gaussian distribution.Kolmogorov-Smirnoff was used for probability density.

---

**Supplementary 1.**
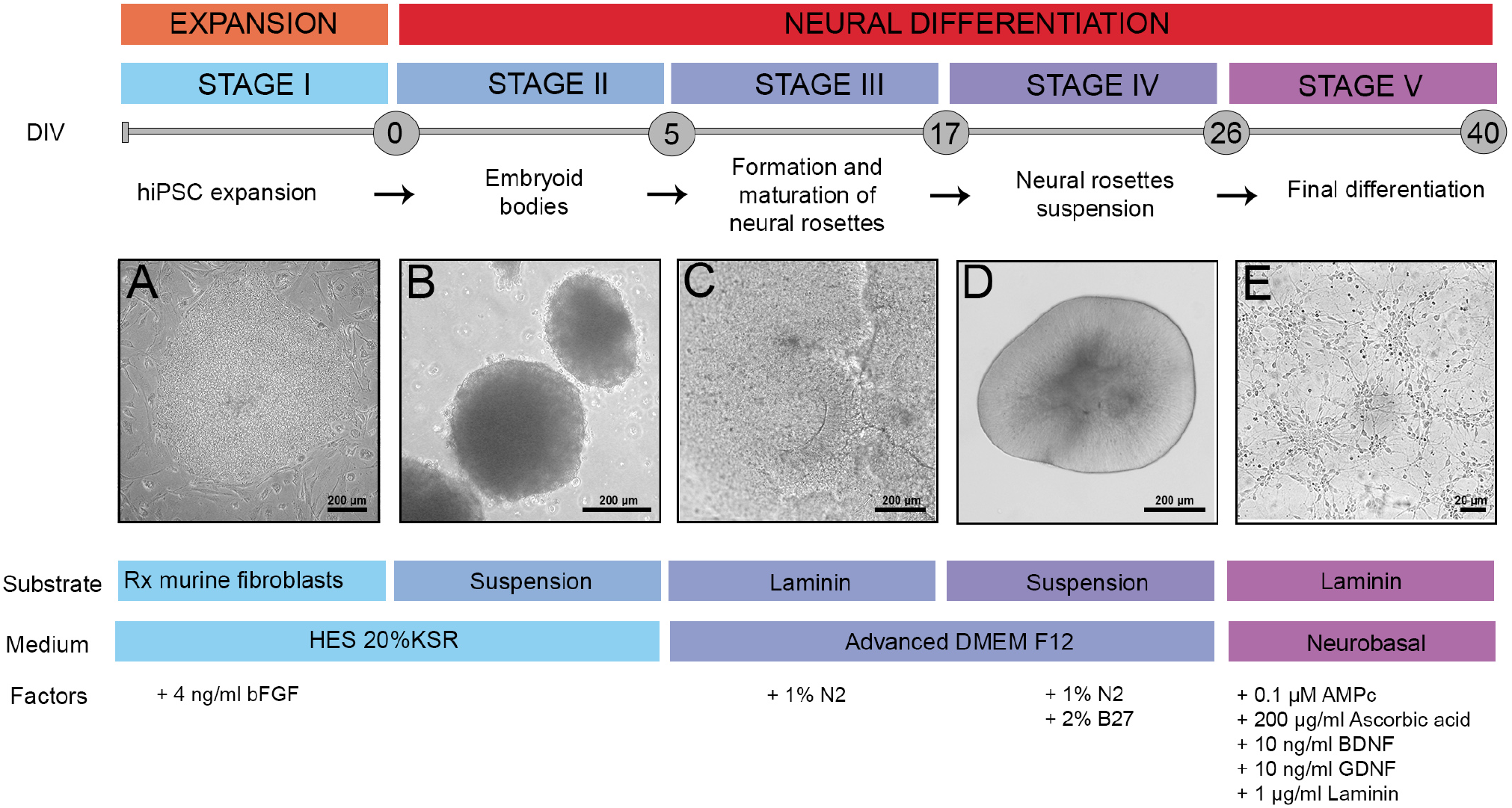
Differentiation of hiPSC to human neurons. (A-E) Brigthfield images of stages in neuronal differentiation from human induced pluripotent stem cells (hiPSC). (A) Colonies of hiPSC from Craig Venter line (CV-hiPSC) on an inactivated mouse embryonic fibroblast (iMEF) feeder layer before enzymatic detachatment. (B) Embrioyd bodies (EBs) in suspension 24 hrs after detachatment. (C) Neural tube-like rosettes plated into laminin-coated well for 7 days. (D) Neural rosettes enriched in neural precursors after mechanic detatachment grown in suspension for 7 days. (E) Plated human neurons 14 days after final differentiation

**Supplementary 2.**
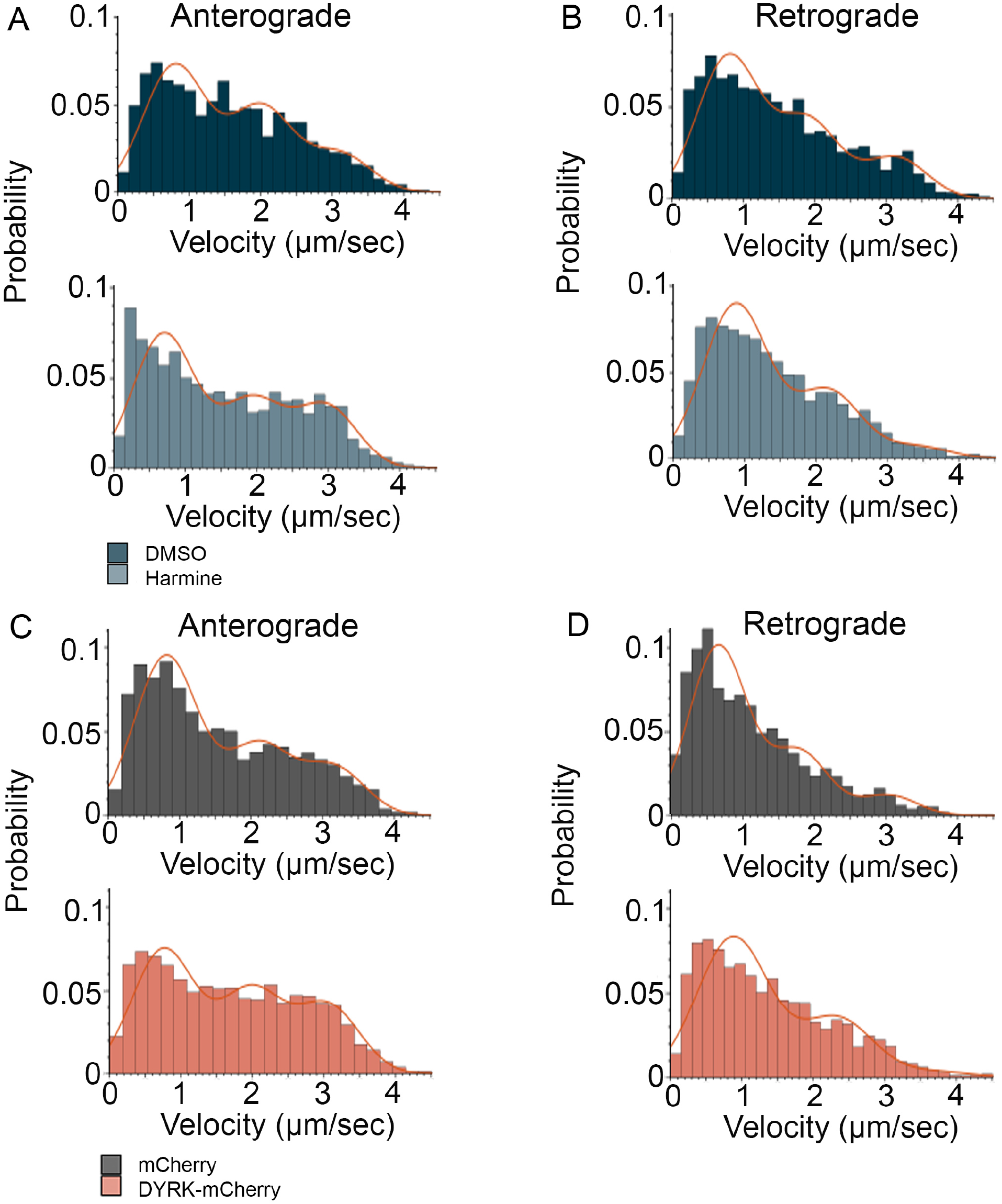
Molecular mechanisms of APP retrograde transport are modulated by harmine and DYRK1A over-expression. (A-D) Histograms showing the distribution of segmental velocities of moving APP-YFP vesicles. Anterograde (A) and retrograde (B) segmental velocities distribution of APP vesicle transport in DMSO (dark blue) and harmine-treated (light blue) human neurons. Anterograde (C) and retrograde (D) segmental velocities distribution of APP vesicle transport in control (APP-YFP+mCherry, grey) and DYRK1A over-expressing (APP-YFP+DYRK-mCherry, red) human neurons. (A-D) Red line represents Gaussian mixture model with three modes associated with number of active motors. Bins width determined by Freedman-Diaconis rule of control condition. Segmental velocities n=7694 anterograde and 6412 retrograde DMSO; 9225 anterograde and 7177 retrograde harmine; 5954 anterograde and 3177 retrograde mCherry; 5788 anterograde and 3762 DYRK-mCherry.

